# Glutamine antagonist DRP-104 suppresses tumor growth and enhances response to checkpoint blockade in *KEAP1* mutant lung cancer

**DOI:** 10.1101/2023.06.27.546750

**Authors:** Ray Pillai, Sarah E. LeBoeuf, Yuan Hao, Connie New, Jenna L. E. Blum, Ali Rashidfarrokhi, Shih Ming Huang, Christian Bahamon, Warren L. Wu, Burcu Karadal-Ferrena, Alberto Herrera, Ellie Ivanova, Michael Cross, Jozef P. Bossowski, Hongyu Ding, Makiko Hayashi, Sahith Rajalingam, Triantafyllia Karakousi, Volkan I. Sayin, Kamal M. Khanna, Kwok-Kin Wong, Robert Wild, Aristotelis Tsirigos, John T. Poirier, Charles M. Rudin, Shawn M. Davidson, Sergei B. Koralov, Thales Papagiannakopoulos

## Abstract

Loss-of-function mutations in *KEAP1* frequently occur in lung cancer and are associated with resistance to standard of care treatment, highlighting the need for the development of targeted therapies. We have previously shown that *KEAP1* mutant tumors have increased glutamine consumption to support the metabolic rewiring associated with NRF2 activation. Here, using patient-derived xenograft models and antigenic orthotopic lung cancer models, we show that the novel glutamine antagonist DRP-104 impairs the growth of *KEAP1* mutant tumors. We find that DRP-104 suppresses *KEAP1* mutant tumor growth by inhibiting glutamine-dependent nucleotide synthesis and promoting anti-tumor CD4 and CD8 T cell responses. Using multimodal single-cell sequencing and *ex vivo* functional assays, we discover that DRP-104 reverses T cell exhaustion and enhances the function of CD4 and CD8 T cells culminating in an improved response to anti-PD1 therapy. Our pre-clinical findings provide compelling evidence that DRP-104, currently in phase 1 clinical trials, offers a promising therapeutic approach for treating patients with *KEAP1* mutant lung cancer. Furthermore, we demonstrate that by combining DRP-104 with checkpoint inhibition, we can achieve suppression of tumor intrinsic metabolism and augmentation of anti-tumor T cell responses.

## Main

Somatic mutations found in cancers play an important role in promoting tumorigenesis by driving multiple hallmarks of cancer including metabolic rewiring and immune evasion^1–4^. As a result, precision medicine-based therapies that directly target driver mutations or downstream dependencies have shown great promise^5–10^. Loss-of-function Kelch-like ECH-associated protein 1 (*KEAP1)* mutations or gain-of-function nuclear factor erythroid 2-related factor 2 (*NFE2L2*, also known as *NRF2)* mutations are found in approximately 20% of lung adenocarcinoma (LUAD) and ∼ 30% of lung squamous cell carcinoma (LUSC)^11, 12^. LUAD and LUSC are the two major histologic subtypes of non-small cell lung cancer (NSCLC). KEAP1 is a negative regulator of NRF2^13–17^, a key transcription factor that governs the cell’s antioxidant response^11, 12, 14, 17^. In LUAD, *KEAP1* mutant tumors respond poorly to checkpoint blockade^18–20^ and are more resistant to KRAS^G12C^ inhibitors ^10, 21, 22^. Unfortunately, there are no clinically approved therapies that specifically target *KEAP1* mutant LUAD.

Multiple pre-clinical studies have demonstrated that *KEAP1* loss leads to NRF2 activation, which promotes LUAD progression and metastasis^2, 3, 23–35^. Our group previously demonstrated that NRF2 activation by *KEAP1* loss rewires cellular metabolism, promotes addiction to extracellular glutamine, and generates a vulnerability that can be targeted by glutaminase (GLS1) inhibition with CB-839^2, 36^. Furthermore, additional studies demonstrated that NRF2 activation in multiple cancers generates a glutamine dependency^2, 37^. However, CB-839 showed limited efficacy in a clinical trial that enrolled patients with *KEAP1* mutant lung cancer possibly because this compound targets only one of many glutamine-dependent reactions that are essential for cancer growth. Therefore, the development of other therapies to target *KEAP1* mutant NSCLC remains a pressing clinical issue.

6-Diazo-5-oxo-L-norleucine (DON), a glutamine antagonist, previously showed promising anti-tumor effects^38^. However, clinical utility was limited due to significant adverse effects^39–42^. Recently DRP-104 (sirpiglenastat), a pro-drug of DON, was developed as a novel cancer agent with reduced toxicity as its activation is dependent on two enzymatic reactions occurring in the tumor microenvironment^43, 44^. However, prior studies evaluating the efficacy of DRP-104 were restricted to subcutaneous tumor mouse models without defined tumor genetics^43, 44^, and conducted in the absence of the native lung microenvironment where anti-tumor immune responses and therapeutic responses can be drastically different^45^. Based on our earlier work^2, 46^, we hypothesize that *KEAP1* mutant lung tumors would be highly sensitive to DRP-104 due to the increased glutamine dependency of these tumors. In this study, we are the first to utilize an antigenic orthotopic lung cancer mouse model and patient derived xenografts (PDXs) to investigate the impact of DRP-104 on *KEAP1* mutant lung tumor growth. We found that in these pre-clinical models, *KEAP1* mutant tumors are highly sensitive to DRP-104, as compared to *KEAP1* wildtype tumors. Using a systematic metabolomics approach, we found that the cell intrinsic sensitivity of *KEAP1* mutant tumors to DRP-104 is primarily mediated by inhibition of nucleotide synthesis. Furthermore, after a comprehensive immune analysis using flow cytometry and multimodal single-cell sequencing (Expanded Cellular Indexing of Transcriptomes and Epitopes by sequencing, also known as ExCITE-seq) in our orthotopic mouse model, we found that DRP-104 reduces exhausted CD4 and CD8 T cell populations, enhances T cell cytokine production, and augments the response to anti-PD1 checkpoint inhibitor therapy in *Keap1* mutant tumors. In summary, our research establishes a convincing mechanistic rationale for the ongoing clinical trial combining DRP-104 with checkpoint blockade in patients with *KEAP1* mutant LUAD (NCT04471415).

## Results

### *KEAP1* mutant tumor growth is impaired by DRP-104

We first sought to investigate the efficacy of the new pro-drug DRP-104 (Fig. 1a) in multiple *KEAP1* mutant and wildtype NSCLC pre-clinical tumor models. To determine the effect of DRP-104 on *Keap1* mutant tumors, we transplanted murine *Kras*^G12D/+^ *p53*^-/-^ *Keap1* knockout (KPK) cell lines generated by CRISPR/Cas9 editing or *Keap1* wildtype (KP) cell lines subcutaneously into immunodeficient mice. We observed that KPK tumors were sensitive to escalating doses of DRP-104, while KP tumors were resistant (Fig. 1b). Interestingly, both cell lines were sensitive to DRP-104 and DON *in vitro* (Extended Data Fig. 1a). Since DON failed in clinical trials due to its high toxicity when delivered systemically, we monitored mice for adverse effects during treatment with multiple doses of DRP-104 and observed no evidence of weight loss or toxicity (Extended Data Fig. 1b). Our previous work has demonstrated that loss-of-function of *Keap1* increases NRF2 transcriptional activity and promotes glutamine addiction in LUAD mouse models^2, 36^. To determine whether sensitivity of KPK cells to DRP-104 is due to this NRF2 mediated glutamine addiction, we overexpressed a gain-of-function mutant of *Nrf2*, which has a deletion in the Neh2 domain and cannot bind KEAP1^2^, in KP cells and measured the sensitivity of this cell line to DRP-104 *in vivo*. Consistent with the loss-of-function *Keap1* mutant tumors, we observed that *Nrf2* gain-of-function tumors were also sensitive to DRP-104 (Fig. 1c). This finding demonstrates that NRF2 activation, and likely the subsequent glutamine addiction it induces, sensitizes cells to DRP-104.

**Figure 1:**
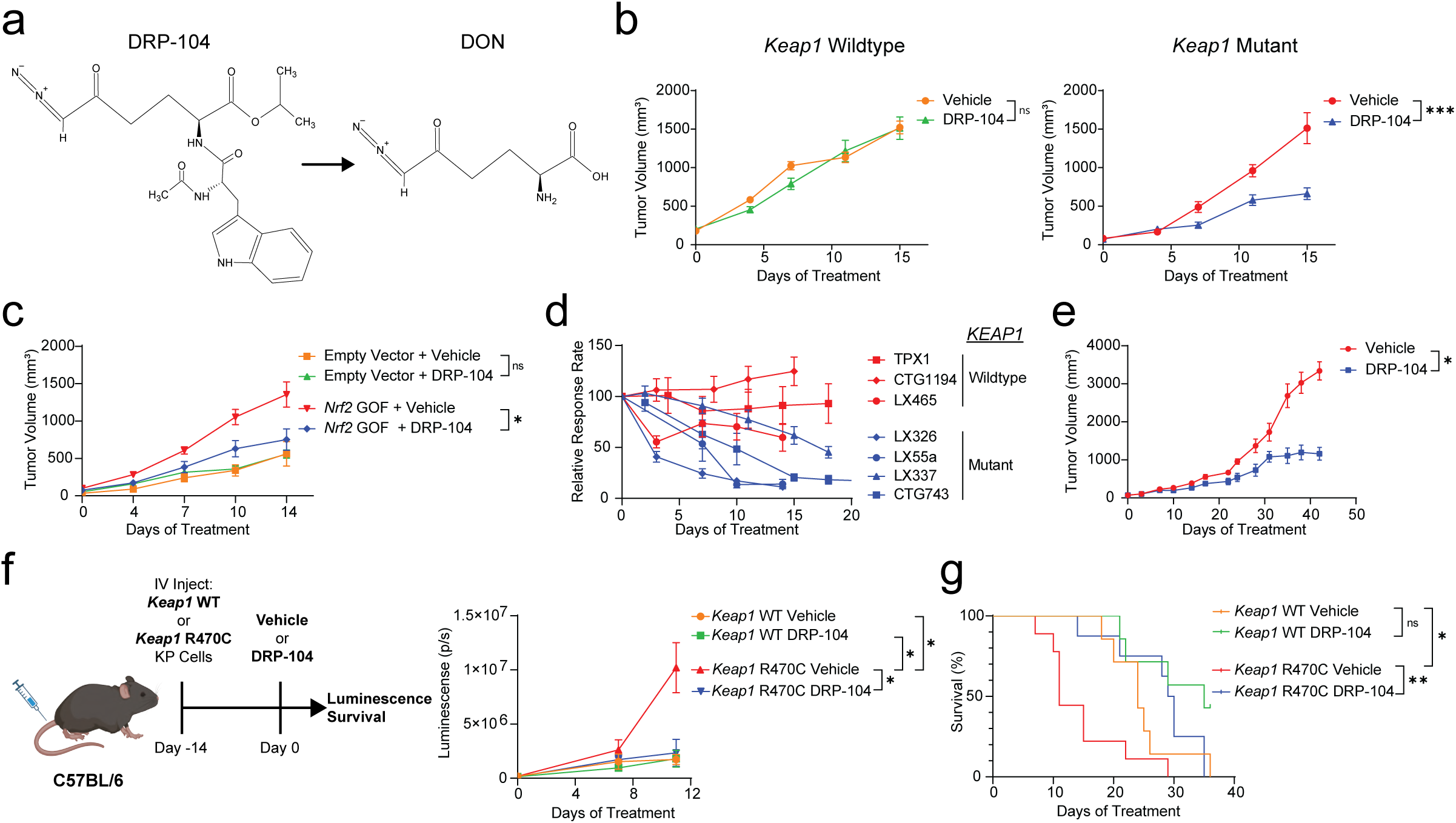
*KEAP1* mutant tumors are sensitive to DRP-104 *in vivo*. **a**: Structure of the prodrug DRP-104 which is enzymatically activated to 6-Diazo-6-oxo-l-norleucine (DON). **b**: *Keap1* wildtype or *Keap1* mutant *Kras*^G12D/+^ *p53*^-/-^ (KP) cells were subcutaneously transplanted into nude mice (n=15 per group). Mice were treated with either DRP-104 (2 mg/kg) or vehicle subcutaneously five days on, two days off. **c**: KP cells were transduced with *Nrf2* gain-of-function (GOF) vector or empty vector control and were subcutaneously transplanted into nude mice (n=7-8 per group). Tumor burden was measured after treatment with either DRP-104 (3 mg/kg) or vehicle control. **d**: Lung adenocarcinoma patient derived xenografts (PDXs) were implanted into NSG mice. Mice were treated with either vehicle control or DRP-104 (3 mg/kg) and tumors were measured over time. Relative response rate (tumor volume/average vehicle volume x 100%) over time is plotted. *KEAP1* wildtype and mutant PDXs are labeled. **e**: Growth kinetics of the lung squamous cell carcinoma PDX LX640 treated with DRP-104 (3 mg/kg) or vehicle control (n=8 per group). **f, g**: Schematic of orthotopic transplant lung cancer model. *Keap1* WT or *Keap1* R470C mutant KP cell lines expressing luciferase were intravenously (IV) injected into C57BL/6 mice on day 0. On day 14 lung luminescence was measured and mice are randomized into treatment groups (7 – 9 mice per group) or either DRP-104 (3 mg/kg) or vehicle control. Tumor growth kinetics based on luminescence as a measure of tumor burden (**f**) and survival data (**g**) is shown. All data is plotted as mean with standard error of the mean. For statistical analysis 2-way ANOVA was used for growth kinetics and Logrank test was used for survival. ns: not significant, * p < 0.05, ** p<0.01, *** p < 0.001, **** p < 0.0001.

*KEAP1* mutations frequently co-occur with serine/threonine kinase 11 (*STK11*, also known as liver kinase B1/*LKB1*) mutations in human LUAD and the co-occurrence of these mutations is associated with resistance to therapies through unknown mechanisms^19^. To verify that the sensitivity of KPK tumors to DRP-104 is retained with loss-of-function mutations in *Stk11*, we generated KPK cells with loss of *Stk11.* When transplanted subcutaneously, these *Keap1/Stk11* mutant tumors were indeed also sensitive to DRP-104, thereby demonstrating that *Stk11* mutations do not induce resistance to DRP-104 (Extended Data Fig. 1c).

Given the substantial degree of genetic heterogeneity in patient tumors, which may lead to drug resistance, we wanted to ascertain the effectiveness of DRP-104 in multiple genetically defined PDX models of NSCLC. To accomplish this, we tested seven LUAD PDX lines (3 *KEAP1* wildtype and 4 KEAP*1* mutant), each with different co-occurring mutations (Extended Data Fig. 2a). *KEAP1* wildtype PDX models did not show a significant response to DRP-104 (TPX1, CTG1194, LX465) (Fig. 1d and Extended Data Fig. 2b). Remarkably, all four *KEAP1* mutant PDXs (LX326, LX55a, LX337, and CTG743) demonstrated a robust response to DRP-104 (Fig. 1d and Extended Data Fig. 2c). Furthermore, DRP-104 maintained long-term suppression of the *KEAP1* mutant PDX CTG743, as demonstrated by significant tumor regression followed by sustained maintenance of tumor growth inhibition during an extended dosing period of fifty days without evidence of resistance (Extended Data Fig. 2c). Interestingly, withdrawal of the drug results in resumption of tumor growth in this PDX suggesting that, in an immunodeficient host, sustained drug administration is required to maintain efficacy (Extended Data Fig. 2c). Since NRF2 activation by either loss-of-function mutation of *KEAP1* or gain-of-function mutation of NRF2 are observed in approximately 30% of lung squamous cell carcinoma (LUSC), the other major subtype of NSCLC^11^, we also tested the therapeutic efficacy of DRP-104 in a LUSC PDX model (LX640) with *KEAP1* mutation. Consistent with the LUAD PDXs, we observed that DRP-104 suppressed the growth of *KEAP1* mutant LUSC (Fig. 1e).

**Figure 2:**
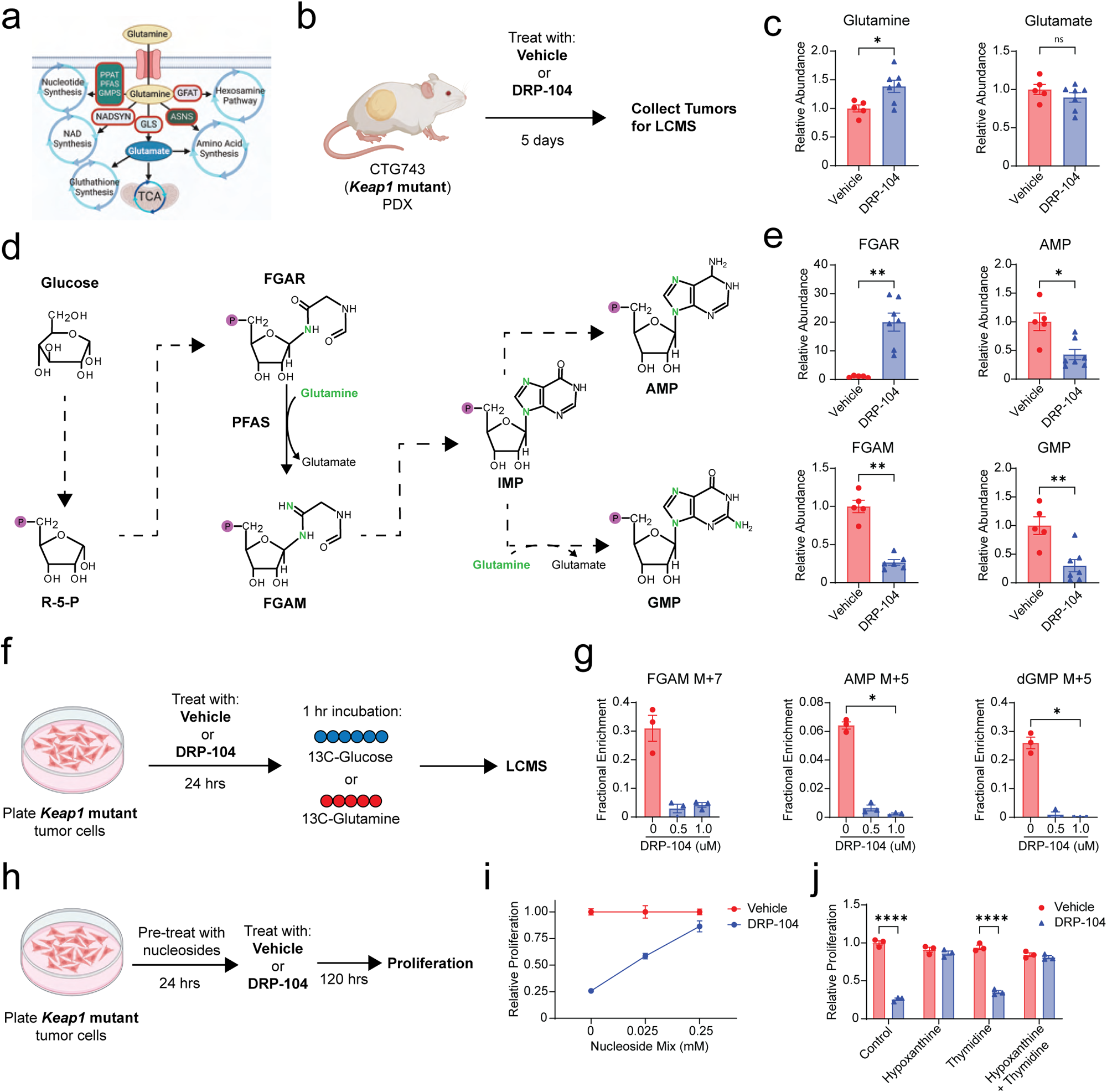
DRP-104 targets nucleotide metabolism in *KEAP1* mutant tumors. **a**: Overview of glutamine dependent synthetic pathways. **b:** Schematic of *in vivo* metabolomics. After implantation of the CTG743 (*Keap1* mutant) patient derived xenograft (PDX) into NSG mice, treatment with DRP-104 (3 mg/kg) or vehicle control (n=5-7 per group) was initiated. After five days of treatment tumors were collected for liquid chromatography mass spectrometry (LCMS). **c**: Relative abundance of glutamine and glutamate as measured by LCMS from experiment in (b). **d**: Diagram outlining key purine biosynthetic steps where glucose is utilized to generate ribose-5-phopshate (R-5-P), FGAR, FGAM, inosine monophosphate (IMP), AMP, and GMP. Nitrogen derived from glutamine is highlighted with a green N. **e**: Relative abundance of FGAR, FGAM, AMP and GMP as measured by LCMS. **f**: Schematic of *in vitro* 13C-Glucose and 13C-Glutamine tracing experiment. *Keap1* mutant tumor cells were treated with DRP-104 (0, 0.5, or 1 uM) (n=3 per group). After 24 hours cells were incubated with either labeled 13C-glucose or 13C-glutamine for 1 hour, then cells were collected for LCMS. **g**: Fractional enrichment of FGAM, AMP, and dGMP for experiment outlined in (f). **h**: Schematic of nucleoside rescue experiment. *Keap1* mutant tumor cells were pre-treated with the nucleosides cytidine, hypoxanthine, uridine, thymidine, guanosine, and adenosine (0 - 0.25 mM) for 24 hours then treated with DRP-104 (2 uM) or control media for 120 hours (n=3 per group). Proliferation was measured by crystal violet. **i**: Plot of nucleoside mix concentration versus relative proliferation of *Keap1* mutant tumor cells treated with DRP-104 normalized to control cells. **j**: Relative proliferation of DRP-104 or vehicle treated *Keap1* mutant tumor cells after addition of either hypoxanthine or thymidine or both. Statistical analysis was done by either Mann Whitney test, Kruskal-Wallis test with Dunn’s multiple comparisons test, or 2-way ANOVA. ns: not significant, * p < 0.05, ** p<0.01, *** p < 0.001, **** p < 0.0001.

Prior work has demonstrated that treatment responses in the lung can markedly differ from those in subcutaneous tissue, a discrepancy partly attributable to anti-tumor immune responses^45^. In addition, inhibiting glutamine metabolism can have profound effects on immune cell function^47, 48^. With this in mind, we sought to evaluate the efficacy of DRP-104 using an antigenic orthotopic lung transplant model that we have recently established^49^. With this model we have demonstrated that KP tumor cell lines expressing a *Keap1* loss-of-function point mutation (R470C) grow faster in the lung than those expressing wildtype *Keap1*, primarily by suppressing CD8 T cell anti-tumor surveillance^49^. Further employing this model, we transplanted *Keap1* R470C mutant or *Keap1* wildtype KP tumor cells, each expressing luciferase, and continuously tracked the lung tumor burden through bioluminescence imaging (Fig. 1f). Upon engraftment of tumors in the lung, mice were treated with DRP-104 or vehicle. Consistent with our subcutaneous *in vivo* model (Fig. 1b), DRP-104 impaired the growth of *Keap1* R470C mutant lung tumors (Fig. 1f) as well as significantly increased the median survival of mice with *Keap1* R470C mutant lung tumors from 11 days to 35 days (Fig. 1g). Overall, using both human and murine tumor models in immunodeficient and immunocompetent mice, we demonstrate that DRP-104 effectively inhibits the growth of *KEAP1* mutant lung tumors, and in some cases, results in significant tumor regression.

### DRP-104 impairs tumor proliferation by inhibition of nucleotide synthesis

Glutamine is used in multiple biosynthetic pathways including nucleotide synthesis, NAD synthesis, glutathione production, hexosamine pathway, and amino acid synthesis, as well as for replenishing TCA intermediates via α-ketoglutarate (Fig. 2a)^50^. We hypothesized that DRP-104 has superior efficacy against *KEAP1* mutant tumors compared to prior selective glutaminase inhibitors due to its ability to target multiple glutamine-dependent metabolic pathways. To probe the metabolic pathways impacted by DRP-104, we performed *in vivo* metabolomics utilizing our *KEAP1* mutant (CTG743) PDX model and harvested tumors after five days of treatment with DRP-104 or vehicle (Fig. 2b). We have previously observed that CB-839 primarily impacted *KEAP1* mutant tumors through reduction of intracellular glutamate via inhibition of GLS1^36^. However, our liquid chromatography mass spectrometry (LCMS) analysis demonstrated that while DRP-104 levels did increase glutamine levels, DRP-104 treatment did not significantly reduce glutamate levels, suggestive that inhibition of glutaminolysis is not a major effect of DRP-104 *in vivo* (Fig. 2c) and acts through mechanisms distinct from CB-839.

We then systematically investigated other metabolic pathways utilizing glutamine (Fig. 2a) to identify the vulnerability of *KEAP1* mutant tumors to DRP-104. Given that nucleotide synthesis plays a pivotal role in cell proliferation, we next focused on the synthesis of purines and pyrimidines. For purine synthesis, the generation of inosine monophosphate (IMP), the purine precursor for AMP and GMP, requires glutamine as a nitrogen source^51^ to generate formylglycinamidine ribonucleotide (FGAM) from formylglycinamide ribonucleotide (FGAR) and this reaction is mediated by the enzyme phosphoribosylformylglycinamidine synthase (PFAS) (Fig. 2d). We found that FGAR abundance was significantly increased while FGAM was significantly reduced in tumors upon DRP-104 treatment, suggesting that the activity of PFAS was inhibited (Fig. 2e). Consistent with an inhibition of the purine synthesis pathway, we found that AMP and GMP levels were both significantly reduced following DRP-104 treatment (Fig. 2e). We then examined pyrimidine biosynthesis, which utilizes glutamine as a substrate for synthesis of the pyrimidine ring (Extended Data Fig. 3a). Orotate, an intermediate in pyrimidine synthesis, and downstream metabolites UMP and dTMP, were significantly reduced following DRP-104 treatment (Extended Data Fig. 3b). Interestingly, CTP was not reduced after treatment with DRP-104 (Extended Data Fig. 3b), despite requiring glutamine for the generation of CTP from UTP.

**Figure 3:**
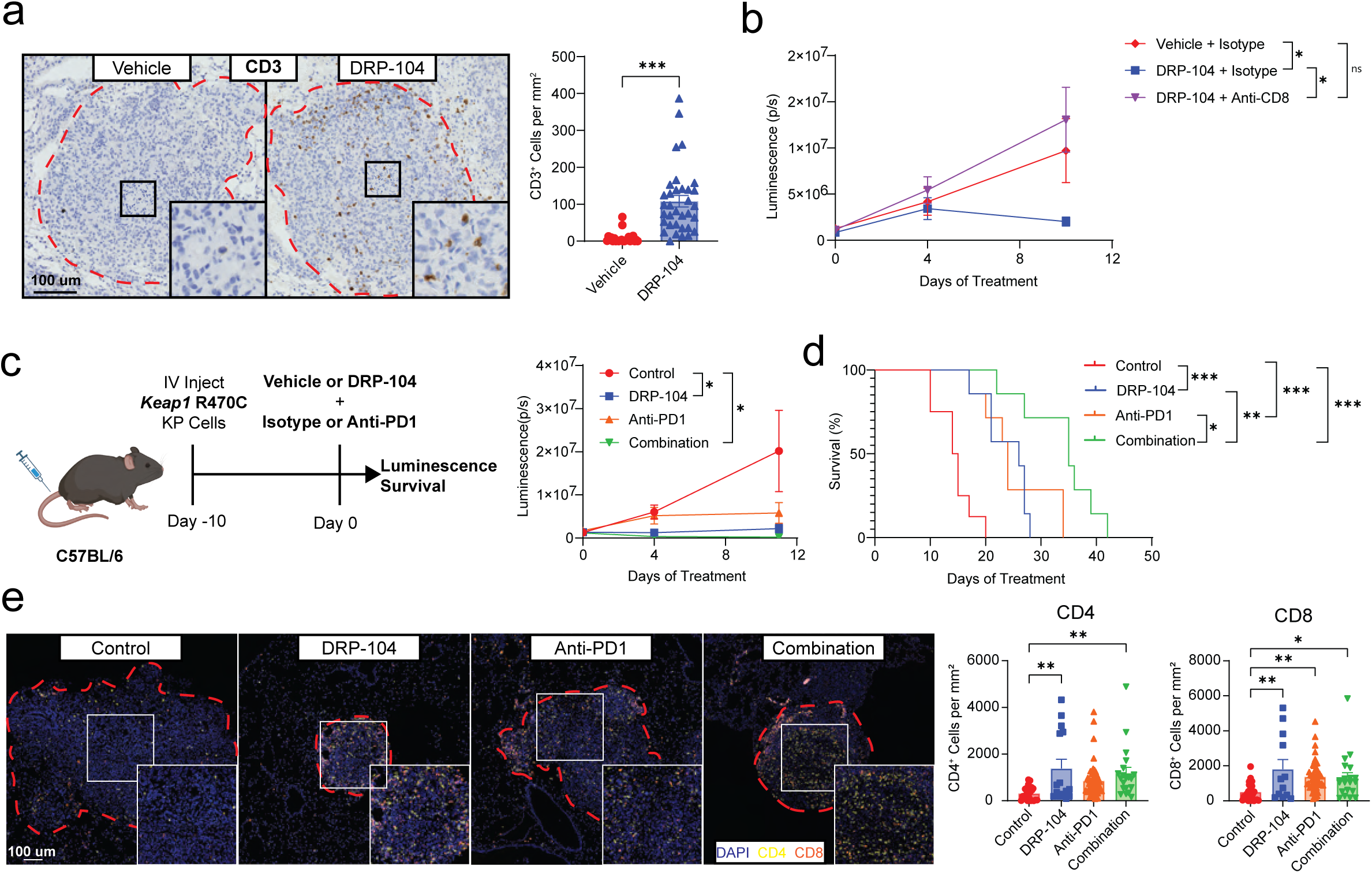
DRP-104 augments T cell infiltration and increase response rates to anti-PD1 therapy. **a**: Immunohistochemistry staining for CD3 of mouse lungs with *Keap1* R470C KP tumors treated with either DRP-104 or vehicle control. Intra-tumoral CD3 quantification is shown for individual tumors. **b**: *Keap1* R470C mutant KP tumor cells were intravenously (IV) injected into C57BL/6 mice. After ten days mice were treated with either anti-CD8 or isotype control (150 ug intraperitoneal twice a week) and DRP-104 or vehicle control (n=4-6 per group). Lung tumor burden as measured by luminescence is displayed. **c**: *Keap1* R470C mutant KP tumor cells were IV injected into C57BL/6 mice. After ten days mice were randomized into treatment conditions displayed in the schematic. Tumor burden was measured over time by luminescence (n=5 per group). **d**, Survival of mice shown from experiment outlined in c. **e**, Multi-color immunofluorescent staining of *Keap1* R470C lung tumors after five days of treatment with DRP-104 (+ isotype), anti-PD1 (+ vehicle), the combination of both, or controls (vehicle + isotype control). Quantification of CD4 (yellow) and CD8 (red) intra-tumoral T cell populations is shown for individual tumors. Data was analyzed by one-way ANOVA and Tukey’s multiple comparison testing or Logrank test. * p < 0.05, ** p<0.01, *** p < 0.001, **** p < 0.0001.

We next examined other glutamine dependent pathways beyond nucleotide synthesis. Glutamine is used as a substrate in the hexosamine pathway in order to synthesize UDP N-acetyl-glucosamine and is also required for the generation of asparagine, aspartate, and glutathione (Fig. 2a). Our LCMS analysis demonstrated that UDP-N-acetyl-Glucosamine and asparagine levels were unchanged, but aspartate and glutathione were significantly reduced by treatment with DRP-104 (Extended Data Fig. 3c).

To identify which synthetic reactions were affected by DRP-104, we performed *in vitro* tracing by labeling DRP-104 treated KPK tumor cells with either 13C-Glutamine or 13C-Glucose for 1 hour (Fig. 2f). *KEAP1* mutant tumors replenish intracellular glutamate via glutaminolysis mediated by the enzyme GLS1 (Extended Data Fig. 3d). Our tracing analysis demonstrated that DRP-104 did not have any effect on glutamate M+5 fractional labeling by 13C-Glutamine (Extended Data Fig. 3d), further suggesting that DRP-104 does not significantly affect glutaminolysis. We chose to focus on the labeling of nucleotide synthesis intermediates given the alteration in purine and pyrimidine synthesis intermediates induced by DRP-104 (Fig. 2e, Extended Data Fig. 3b). First, we looked at labeling of intermediates of purine synthesis by 13C-Glucose (Extended Data Fig. 3e) and found a trend towards a reduction in the FGAM M+7 fraction (Fig. 2g). In addition, there was a decrease in labeled AMP M+5 and dGMP M+5 fraction (Fig. 2g), demonstrating that DRP-104 impairs the biosynthetic reactions for these metabolites. Similarly, when examining labeled pyrimidines, we found that DRP-104 reduced the labeling of UMP, dTMP, and CTP (Extended Data Fig. 3f), suggesting that DRP-104 also impairs the synthesis of pyrimidines.

To determine whether impaired nucleotide synthesis was indeed responsible for the reduction in tumor growth induced by DRP-104, we investigated whether the addition of nucleosides could restore the proliferation in treated *Keap1* mutant tumor cells. We pretreated KPK tumor cells *in vitro* with a mix of nucleosides (cytidine, thymidine, hypoxanthine, uridine, guanosine, and adenosine). After 24 hours of nucleoside pre-treatment, we administered DRP-104 to the KPK cells and measured their proliferation five days later (Fig. 2h). Remarkably, the proliferation of DRP-104 treated KPK cells improved drastically in a dose-dependent manner with increasing nucleoside concentrations (Fig. 2i). To distinguish the necessity of purines from pyrimidines in DRP-104 treated cells, we pre-treated KPK cells *in vitro* with either hypoxanthine, thymidine, or both. Addition of the purine hypoxanthine, but not the pyrimidine thymidine, rescued the proliferation of DRP-104 treated *Keap1* mutant tumor cells (Fig. 2j) demonstrating that while both purine and pyrimidine synthesis pathways are inhibited with DRP-104, the deficit in purines drives the reduced proliferation observed.

To comprehensively evaluate the role of other metabolic deficiencies induced by DRP-104 we performed additional metabolite rescues. While loss-of-function mutations in *KEAP1* result in increased expression of antioxidant pathways, our previous work has established that the sensitivity to glutaminase inhibition is not due to increased oxidative stress^36^. Rather, CB-839 suppresses tumor growth by reducing intracellular glutamate stores required for TCA cycle anaplerosis^36, 52^ and amino acid synthesis - a suppression reversible by either glutamate supplementation or blocking glutamate export through the transporter xCT using erastin^36^. Contrary to the effect seen with CB-839, addition of glutamate or erastin had no impact on cell proliferation of DRP-104 treated KPK cells (Extended Data Fig. 3g) demonstrating that DRP-104’s effect is not mediated through intracellular glutamate depletion. Supplementation with cell permeable α-ketoglutarate (DMG) or pyruvate also did not improve cell proliferation, suggesting that DRP-104 does not induce deficiencies in TCA cycle intermediates (Extended Data Fig. 3g). Finally, treatment with the aspartate or the antioxidant Trolox also failed to rescue DRP-104 treated cells despite having reduced aspartate and glutathione levels (Extended Data Figs. 3c and 3g). We therefore concluded that while DRP-104 induces multiple metabolic deficiencies, its primary mechanism of impairing the proliferation of *KEAP1* mutant tumors is through inhibition of purine synthesis.

### DRP-104 modulates anti-tumor T cell responses and augments checkpoint blockade efficacy

The function of T cells is heavily influenced by nutrient availability. Activated T cells are highly proliferative and utilize numerous metabolites including glucose, asparagine, and serine^53–59^. CD8 T cell function is significantly impaired in glutamine depleted conditions^60, 61^. Additionally, differentiation of CD4 T cells into various subsets, including Th1, Th17, and Tregs, is modulated by the levels of glutamine and glutamate^48^. Furthermore, prior work has suggested that the efficacy of DON *in vivo* is partially mediated by the enhancement of CD8 T cell function^47^. Recently, despite efficacy in pre-clinical models^2, 62, 63^, blockade of GLS1 with CB-839 had failed to show efficacy in clinical trials. One possible explanation is that the beneficial effect of CB-839 on *KEAP1* mutant tumors may be offset by potential negative effects on T cell function^60^. We therefore thought it was necessary to evaluate the impact of DRP-104 on the immune microenvironment of *Keap1* mutant lung tumors. We first sought to determine if T cell infiltration is altered by DRP-104 in *Keap1* mutant lung tumors. To investigate this, we performed immunohistochemistry for CD3 to quantify T cell infiltration of end stage *Keap1* R470C mutant lung tumors. In vehicle treated mice, *Keap1* mutant tumors demonstrated immune exclusion with very low intra-tumoral T cell infiltration (Fig. 3a). However, DRP-104 significantly increased the infiltration of T cells into tumors (Fig. 3a). These findings suggest that one of the mechanisms by which DRP-104 might suppress tumor growth *in vivo* is by enhancing anti-tumor T cell responses against *Keap1* mutant tumors.

We previously showed that CD8 T cell depletion had no impact on *Keap1* mutant tumor growth using this model^49^. To determine whether T cells were now generating anti-tumor responses in the context of the glutamine antagonist, we evaluated the effect of CD8 T cell depletion on DRP-104 treated *Keap1* mutant tumors. To do this, we injected mice with *Keap1* R470C mutant cells via tail vein injection. After tumors had successfully engrafted in the lung, we treated mice with vehicle and isotype control, DRP-104 with isotype control, or DRP-104 with anti-CD8 depleting antibody. In contrast to our work where anti-CD8 antibodies had no effect on *Keap1* mutant tumor growth^49^, CD8 T cell depletion accelerated the growth of *Keap1* R470C mutant tumors in the presence of DRP-104 (Fig. 3b) suggesting that DRP-104 can activate anti-tumor T cell responses against *Keap1* mutant tumors.

Patients with *KEAP1* mutant LUAD are known to have poor responses to checkpoint blockade^19, 64^, and our previous work has similarly demonstrated that this also holds true in our *Keap1* mutant orthotopic tumor mouse model^49^. Given our findings that DRP-104 enhances T cell infiltration and induces T cell mediated anti-tumor responses (Figs. 3a and 3b), we next asked whether DRP-104 can augment anti-tumor responses when combined with standard of care checkpoint blockade. Following the injection of *Keap1* R470C mutant KP cells, we administered either DRP-104 or a vehicle control to the mice, along with either anti-PD1 or isotype control antibody (Fig. 3c). Lung tumor burden was then monitored via bioluminescence. Consistent with previous results, tumor growth was impaired with DRP-104 (Fig. 3c). While anti-PD1 alone dampened tumor growth, combination of anti-PD1 with DRP-104 significantly reduced tumor growth (Fig. 3c) and markedly increased survival of mice with *Keap1* mutant lung tumors (with a median survival difference exceeding 20 days) (Fig. 3d).

Given the increased infiltration of T cells and their functional importance in suppression of tumor growth in response to DRP-104, we next evaluated which T cell subsets were impacted in the tumor microenvironment of animals treated with DRP-104 alone and in combination with anti-PD1. We collected tumor bearing lungs after five days of treatment to minimize differences in immune infiltration associated with disparities in tumor burden. Using multi-immunofluorescence, we stained for CD4 and CD8 T cells and quantified the intra-tumoral populations (Fig. 3e). We found that both CD4 and CD8 populations were increased with DRP-104 treated animals compared to the control group. Anti-PD1 monotherapy resulted in an increase in CD8 infiltration without significantly altering CD4 infiltration. While the combination of DRP-104 and anti-PD1 increased the infiltration of CD4 and CD8 T cells compared to the control arm, this infiltration was not significantly different than the DRP-104 single treatment condition. The fact that combination therapy significantly improved survival compared to monotherapy (Fig. 3d) despite comparable T cell infiltration to the treatment groups (Fig. 3e) raises the possibility that the functionality of T cells is enhanced with the combination of DRP-104 with anti-PD1 rather than simply increasing the number of T cells. Our findings demonstrate that DRP-104 exerts tumor-intrinsic effects by suppressing glutamine-dependent metabolism, while also promoting anti-tumor T cell responses, thereby enhancing the efficacy of checkpoint blockade.

### Multimodal sequencing identifies T cell populations altered by DRP-104

Intra-tumoral T cells are a heterogenous population comprised of effector, exhausted, and memory CD8s, along with various CD4 T cell subsets such as Th1, Th17, and Tregs. Given the general diversity of T cell populations and our observation that DRP-104 may alter the functionality of T cells, we chose to comprehensively examine the immune microenvironment of *Keap1* mutant tumors utilizing the multimodal single-cell platform ExCITE-seq^65, 66^ to identify immune populations that are impacted by glutamine antagonism. ExCITE-seq uses oligo-tagged antibodies to simultaneously provide surface epitope information along with gene expression at a single-cell resolution^65–67^. We treated mice with *Keap1* R470C mutant lung tumors with DRP-104 and/or anti-PD1 and harvested whole lungs for single-cell analysis after five days of treatment (Fig. 4a), a timepoint where tumor burden differences amongst treatment arms are minimal (Fig. 3c). Lungs were digested and non-circulating CD45^+^ immune cells and tumor populations we sorted out and subsequently analyzed by ExCITE-seq (Fig. 4a). Our initial clustering revealed a diverse subset of immune populations including macrophages, neutrophils, B cells, T cells, and NK cells (Fig. 4b). In general, we observed a subtle increase in total T cell populations with combination therapy, further supporting that functional changes may contribute to the robust suppression in tumor growth by CD8 T cells (Fig. 3b). We focused our single-cell analysis on populations comprised of T cells, NKT cells, NK cells, and ILCs with clustering of these populations shown in Fig. 4c. Utilizing antibody derived tags (ADTs) we were able to clearly identify CD4 and CD8 T cell populations (Extended Data Fig. 4a). When looking at relative gene expression in CD4 and CD8 T cells, we observed a clear upregulation of several genes associated with activation, such as *Cd44*, *Pdcd1*, and *Nkg7* brought about by DRP-104 and/or anti-PD1. Concurrently, we noticed a downregulation of genes associated with naïve populations, such as *Ccr7* and *Sell* (Extended Data Fig. 4b)

**Figure 4:**
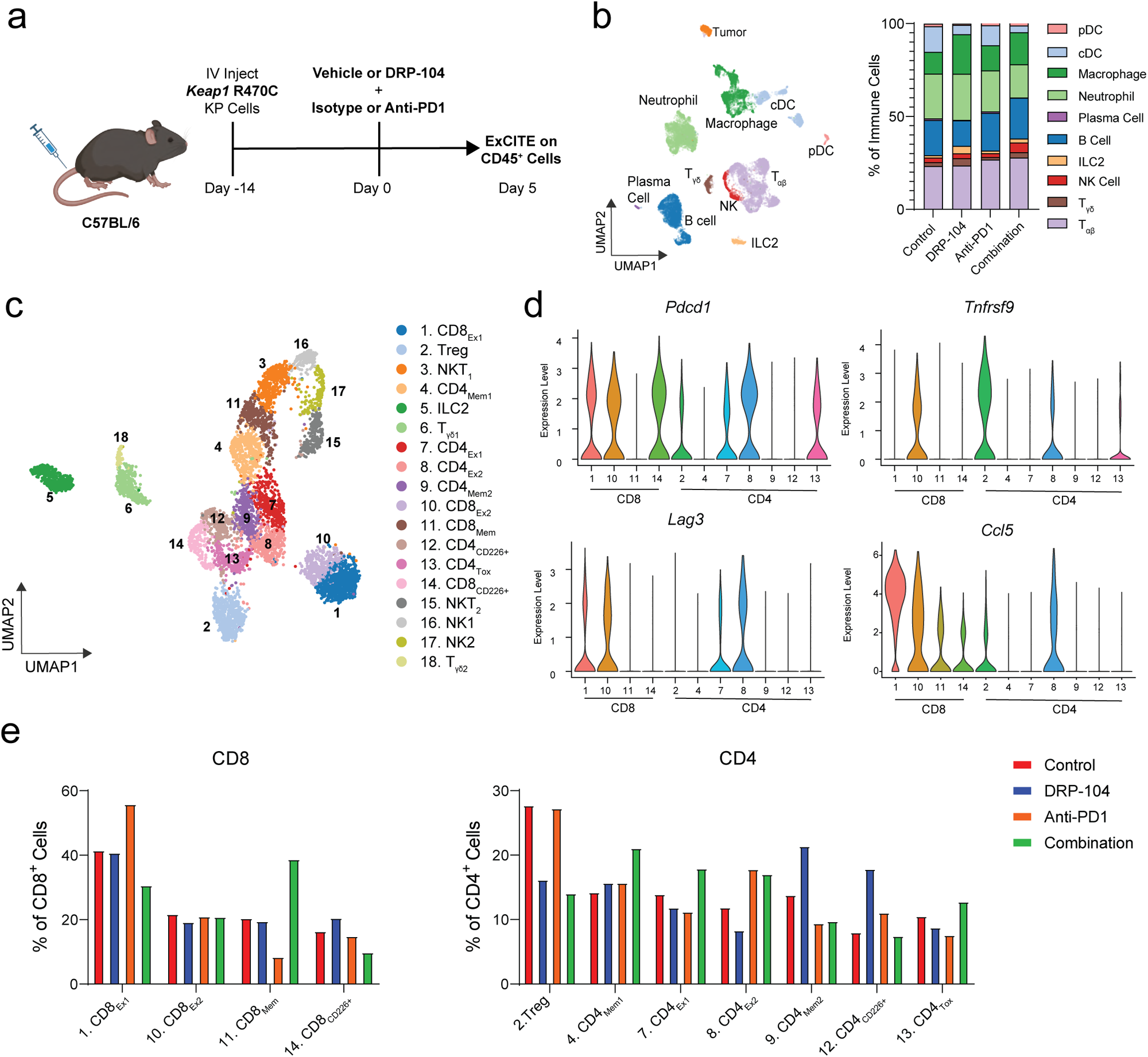
ExCITE-seq identifies transcriptional changes in T cell populations with DRP-104 and anti-PD1 therapy. **a**: Schematic of experimental design of acquisition of samples for ExCITE-seq. Fourteen days after injection of *Keap1* R470C mutant KP cells, mice were treated with DRP-104 (3mg/kg) or vehicle and anti-PD1 (200 ug intraperitoneal every other day) or isotype control. Tumor bearing lungs were digested after five days of treatment and extravascular CD45^+^ cells and tumor cells were sorted for analysis by ExCITE-seq (n=2 per group). **b**: UMAP showing clustering of cell populations with quantification of immune subpopulations. **c**: Subcluster showing T cell, NKT cell, NK cell, and ILC populations with subclusters labelled. **d**: Violin plots showing expression of *Pdcd1*, *Tnfrsf9*, *Lag3*, and *Ccl5* by T cell clusters shown in (c). **e**: Quantification of T cell subclusters stratified by treatment condition and normalized to total CD8 or CD4 T cells.

To gain greater understanding of the diverse adaptive immune populations and transcriptional changes induced by DRP-104, we subclustered the T cell, ILC, NK, and NKT cells into 18 subclusters (Fig. 4c). Seven of these clusters were identified as CD4 T cells, and four as CD8 T cells (Fig. 4c, Extended Data Fig. 4a). Differential gene expression and ADT expression facilitated identification of these T cell populations (Fig. 4d, Extended Data Figs. 4c and 4d). We identified four CD8 T cell clusters with two of these clusters (labeled CD8_Ex1_ and CD8_Ex2_) co-expressing *Lag3* and *Pdcd1* (Fig. 4d), likely representing some degree of exhaustion^68–71^. These two exhausted T cell clusters distinguish themselves with CD8_Ex1_ displaying higher *Ccl5* expression while CD8_Ex2_ expresses *Tnfrsf9* (Fig. 4d), another marker of exhaustion. Notably, *Ccl5* expressing CD8 T cells have previously been implicated with a dysfunctional state and poor responses to anti-PD1 therapy in orthotopic lung cancer models^45^. Our data demonstrates that anti-PD-1 therapy expands these *Ccl5* expressing dysfunctional CD8 T cells (CD8_Ex1_) (Fig. 4e) within the lungs of tumor-burdened mice, potentially limiting the effectiveness of anti-PD1 therapies. However, addition of DRP-104 reduces this population, possibly facilitating more effective anti-tumor responses (Fig. 4e). We also identified a memory CD8 cluster (CD8_Mem_) expressing Ly6C^72^ (Extended Data Fig. 4e) that is preferentially expanded with combination therapy (Fig. 4e).

In a parallel manner, we looked at differences in CD4 subsets induced by DRP-104, anti-PD1, or combination therapy. Quantification of these subclusters revealed that the most dramatically reduced CD4 population by DRP-104 are T regulatory cells (Tregs) (Fig. 4e), which we have previously shown to be enriched in *Keap1* mutant tumors^49^. A similar change in this population was observed upon both DRP-104 treatment and in response to the combination therapy. Similar to CD8 T cells, we identified two CD4 populations with co-expression of *Pdcd1* and *Lag3* expression (shown in Fig. 4d, labeled as CD4_Ex1_ and CD4_Ex2_ in Figs. 4c and 4e), suggestive of an exhausted state. Interestingly, both of these exhausted clusters, along with the *Tox*-expressing CD4_Tox_ cluster, were slightly decreased by DRP-104 treatment, but paradoxically expanded upon combination therapy. However, combination therapy also expanded a CD4 memory population (CD4_Mem1_), similar to memory CD8 T cells (CD8_Mem_) (Fig. 4e). In summary, our single-cell analysis demonstrates that combining DRP-104 with anti-PD1 may alter the functionality and transcriptional state of *Keap1* mutant tumor infiltrating CD4 and CD8 T cell populations driving them from an exhausted program towards a more functional effector/memory state.

### DRP-104 enhances the effector function of CD4 and CD8 T cells

Our ExCITE-seq data revealed that DRP-104 and anti-PD1 therapy had a dramatic effect on specific T cell populations, including Tregs, and on the distribution of memory T cells versus exhausted T cell subsets. We then aimed to corroborate these findings and further characterize the functionality of T cells within the tumor microenvironment of *Keap1* mutant lung tumors treated with DRP-104. We performed multi-color flow cytometry on *Keap1* R470C mutant KP tumor bearing lungs (gating shown in Extended Data Fig. 5a) after five and ten days of treatment with DRP-104, anti-PD1 or the combination of both (Fig. 5a). After five days of treatment, we observed a modest increase in CD3 T cells following combination therapy with anti-PD1 and DRP-104, in comparison to untreated controls (Fig. 5b). Despite not being statistically significant, this increase in T cell population is likely driven by small increases in CD4 and CD8 T cell subsets (Extended Data Fig. 5b). Consistent with our ExCITE-seq data, DRP-104 profoundly decreased the proportion of Tregs (Figs. 4e and 5c). Furthermore, we observed an increase in CD8 central memory populations (identified by expression of CD44^+^ and CD62L^+^) following DRP-104 treatment which was further augmented with addition of anti-PD1 (Fig. 5d).

**Figure 5:**
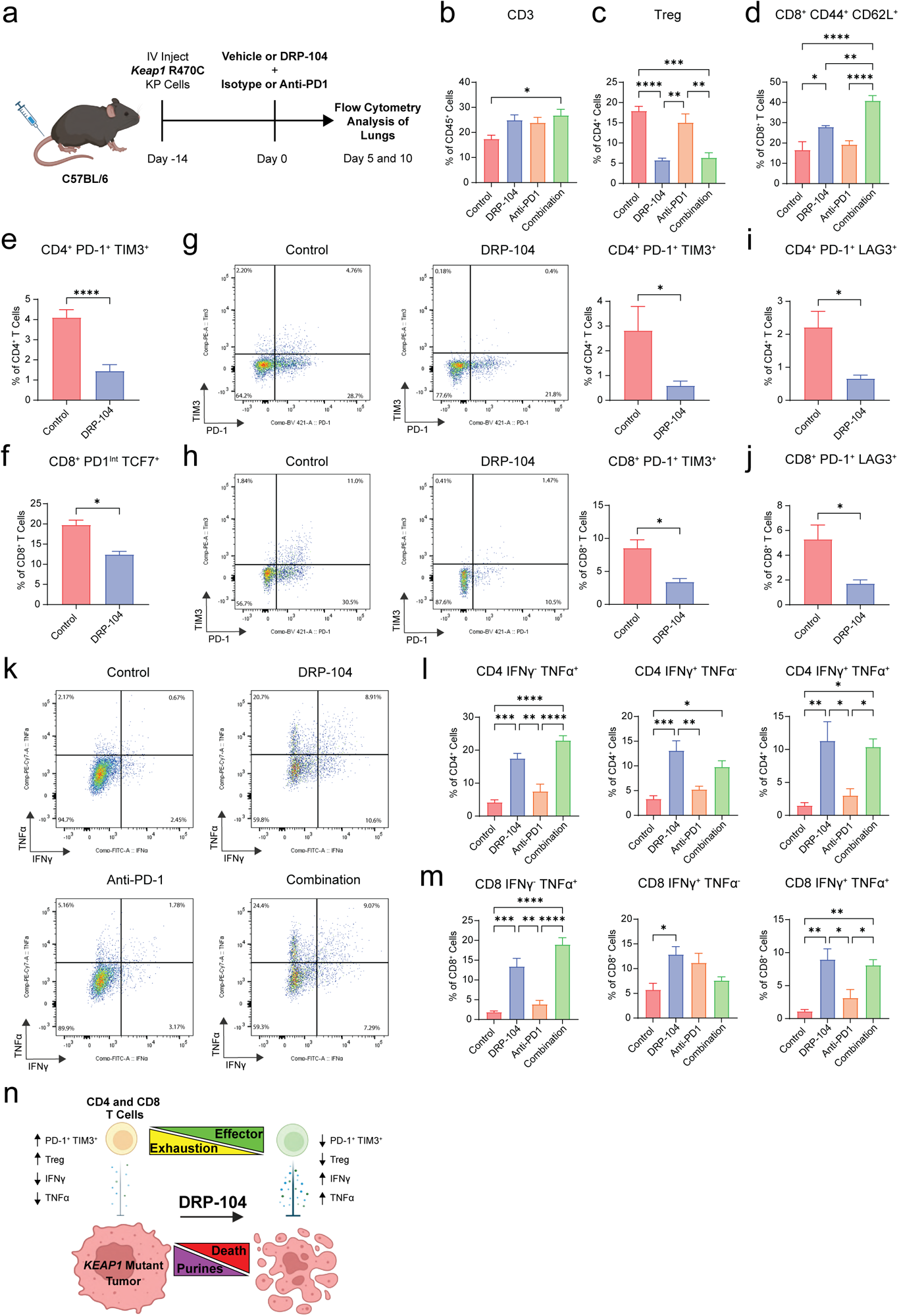
DRP-104 reduces T cell exhaustion and enhances effector T cell function *in vivo*. **a**: Schematic of experimental design. *Keap1* R470C mutant KP lines were injected intravenously into C57BL/6 mice. Fourteen days after injection, treatment with DRP-104 (3 mg/kg) or vehicle and anti-PD1 (200 ug intraperitoneal three times a week) or isotype control was initiated. Lungs were collected from tumor bearing mice either five or ten days after treatment initiation and analyzed by flow cytometry. **b**, **c**, **d, e, f**: Flow cytometry quantification of (**b**) CD3 T cells, (**c**) Tregs (CD4^+^ FoxP3^+^ CD25^+^), and (**d**) CD8^+^ CD44^+^ CD62L^+^ (central memory CD8 T cells), (**e**) CD4^+^ PD-1^+^ TIM3^+^, and (**f**) CD8^+^ PD-1 intermediate TCF7^+^ populations after five days of treatment (n=5 per group). **g, h**: Flow cytometry analysis of PD-1^+^ TIM3^+^ populations for (**g**) CD4 T cells and (**h**) CD8 T cells with representative gating after ten days of treatment with DRP-104 (n=3-6 per group). **i, j:** Flow cytometry quantification of PD-1^+^ LAG3^+^ for (**i**) CD4 T cells and (**j**) CD8 T cells (n=3-6 per group) after ten days of treatment with DRP-104. **k, l, m**: Representative gating (**k**) and flow cytometry quantification of IFNγ and TNFα expression for PMA/ionomycin stimulated (**l**) CD4 T cells and (**m**) CD8 T cells after five days of treatment with DRP-104 or vehicle and/or anti-PD1 or isotype control (n=5 per group). **n**: Overview of effect of DRP-104 on *KEAP1* mutant tumors and T cells. Data was analyzed by either Mann Whitney test or one-way ANOVA and Tukey’s multiple comparison testing. * p < 0.05, ** p<0.01, *** p < 0.001, **** p < 0.0001.

We next shifted our focus on evaluating T cell exhaustion since our ExCITE-seq analysis suggested that DRP-104 reduced T cell exhausted subsets (Fig. 4e). Based on several studies^68, 69, 71, 73–77^, T cell exhaustion is a state of dysfunction associated with expression of surface inhibitor markers such as PD-1, TIM3, TIGIT, and LAG3 with PD-1/TIM3 co-expression denoting terminally exhausted T cell populations^71, 78^. This state is often characterized by reduced functionality, typically manifested by impaired cytokine production^68, 69, 71, 73–76^. Although exhaustion has been most extensively studied in the context of CD8 T cells, similar gene expression profiles are also seen in exhausted CD4 T cells^73^. Flow cytometry analysis revealed that five days of treatment with DRP-104 significantly reduced the proportion of terminally exhausted PD1^+^ TIM3^+^ CD4 T cells, but not TIM3^+^ PD1^+^ CD8 T cells (Fig. 5e, Extended Data Fig. 5c). Unfortunately, likely due to the blocking effect of anti-PD1 therapy, we were unable to identify PD1^+^ populations in mice treated with anti-PD1 therapy. Despite not observing changes in CD8^+^ PD-1^+^ TIM3^+^ T cells, we did find a reduction in PD-1^Int^ TCF7^+^ CD8 T cells in DRP-104 treated animals (Fig. 5f), corresponding to CD8 exhausted progenitor cells^70, 79–81^. These progenitor cells are thought to give rise to terminally exhausted T cells but can also be rescued to differentiate into memory/effector T cells^79, 80, 82^. It is feasible that the increase in CD8 memory cells observed with DRP-104 treatment is due to driving exhausted progenitor CD8 T cells towards memory cell differentiation. Since T cell exhaustion is induced by prolonged antigen stimulation^83^ and because of our observation that DRP-104 treatment reduced exhausted progenitor CD8 T cells, we hypothesized that additional days of DRP-104 treatment would drive a reduction in the proportion of terminally exhausted CD8 T cells. After ten days of DRP-104 treatment, when we quantified terminally exhausted CD4 and CD8 populations, we indeed observed that both terminally exhausted CD4 and CD8 populations co-expressing PD-1 and TIM3 (Figs. 5g and 5h) or co-expressing PD-1 and LAG3 (Figs. 5i and 5j) were significantly reduced.

A fundamental characteristic of T cell exhaustion is reduced T cell function that is typified by decreased production of cytokines^69^, such as IFNγ and TNFα, which are key effector molecules in anti-tumor responses. We have previously demonstrated that these cytokines are suppressed in T cells from *Keap1* mutant tumors compared to wildtype tumors^49^. To assess whether DRP-104 treatment improves the function of both CD4 and CD8 T cells, we isolated T cells from *Keap1* mutant tumors treated with DRP-104 and/or anti-PD1 and evaluated their cytokine production. Flow cytometry analysis revealed that CD4 and CD8 T cells from DRP-104 treated mice had augmented IFNγ and TNFα production (Figs. 5k, 5l and 5m) suggestive that the effector function of these T lymphocytes was greatly enhanced by the drug. Specifically, we find that DRP-104 treatment resulted in increased IFNγ^+^ TNFα^+^ double positive T cells (Figs. 5l and 5m). Interestingly, anti-PD1 alone or the combination of DRP-104 with anti-PD1 did not necessarily increase the expression of these effector cytokines when compared to DRP-104 monotherapy (Figs. 5l and 5m). Using complementary methods, including ExCITE-seq and functional flow cytometry assays, we established that DRP-104 not only targets tumor intrinsic purine metabolism but also diminishes Tregs and exhausted T cell populations that characterize *Keap1* mutant lung tumors. Based on these observations, we conclude that DRP-104 therapy increases the functionality of anti-tumor CD4 and CD8 T cell responses resulting in overall improved outcomes when combined with anti-PD1 therapy (Fig. 5n).

## Discussion

*KEAP1* mutations are frequently found in LUAD and are associated with poor response rates to standard of care therapy^2, 12, 19, 49, 64^. Despite knowing the metabolic vulnerabilities of *KEAP1* mutant LUAD^2, 23, 36^, there are currently no clinically approved treatments specifically targeting this mutation. The KEAPSAKE trial was a phase 2 randomized multicenter double blind clinical trial comparing the addition of the glutaminase inhibitor CB-839 or placebo control to standard of care checkpoint blockade and chemotherapy for patients with metastatic NSCLC with *KEAP1* or *NRF2* mutations. The trial was terminated due to lack of clinical benefit and therefore alternative approaches to targeting *KEAP1* mutant lung tumors need to be explored. Here using immunodeficient and immunocompetent orthotopic murine cancer models we demonstrate that DRP-104, a novel broad acting glutamine antagonist, is efficacious against *KEAP1* mutant tumors by a mechanism distinct from GLS1 selective inhibitors, such as CB-839. Through our metabolomic analysis we show that the intrinsic vulnerability of *KEAP1* mutant tumors to DRP-104 arises from the inhibition of nucleotide synthesis. Utilizing our recently developed antigenic orthotopic lung cancer model, we revealed that not only is DRP-104 effective in targeting *Keap1* mutant lung tumors, but that in combination with checkpoint blockade it led to significantly enhanced survival of mice compared to monotherapy. Using ExCITE-seq, we further discovered that DRP-104 treatment leads to notable reduction in Treg cells. In addition, combining DRP-104 with anti-PD1 reduces a previously described *Ccl5* expressing dysfunctional CD8 population (CD8_Ex1_)^45^, while also expanding memory T cell populations (CD8_Mem_ and CD4_Mem1_). We then validated that DRP-104 reduced exhausted T cell populations and improved the functionality of T cells, as demonstrated by increased IFNγ and TNFα expression. Overall, our work validates that *Keap1* mutant tumors are sensitive to inhibition of glutamine metabolism by DRP-104, which operates both through cell intrinsic mechanisms and also through enhancement of anti-tumor T cell responses.

We identified that the major cell intrinsic mechanism that contributes to the high sensitivity of *KEAP1* mutant tumors to DRP-104 centers on nucleotide synthesis. Previous studies from our lab demonstrated that *KEAP1* mutant tumors are glutamine addicted and depend on exogenous glutamine^2, 36^. However, unlike CB-839 which impairs glutamate-dependent anaplerosis, DRP-104 targets multiple glutamine-dependent reactions in *KEAP1* mutant tumors. Through a comprehensive metabolic analysis, we identified purine synthesis as a major target of DRP-104. Moreover, supplementation with nucleosides proved sufficient to rescue the DRP-104 mediated inhibition of *Keap1* mutant cell growth. A second possible tumor intrinsic mechanism contributing to the sensitivity to DRP-104 is due to the enzymatic activation of the prodrug by the enzyme carboxylesterase CES1 to produce the active form DON. *CES1*, and the mouse ortholog *ces1g*, are transcriptional targets of NRF2^2, 84^. As a result, we speculate that the tumor microenvironment of *KEAP1* mutant tumors could be enriched with DON, the active form of DRP-104. This could potentially lead to a significantly enhanced efficacy of the drug. Additional work is needed to validate role of NRF2 in *Ces1g* expression and DRP-104 sensitivity. CES1 expression could potentially serve as a biomarker in any tumor type that is sensitive to DRP-104.

Targeting glutamine metabolism is a double-edged sword with potential consequences to proliferating cells. Prior work has demonstrated that T cell activation and proliferation is dependent on glutamine^61^. However, glutamine utilization has multiple effects on T cell effector function^47, 60, 61, 85, 86^. Interestingly, it has been suggested that inhibition of GLS1, the enzyme that metabolizes glutamine to glutamate, with CB-839 can either enhance or impair CD8 cytotoxic function^60, 85^. Other work has demonstrated that GLS1 inhibition can enhance Th1 cytokine production^48^. The work presented here does not clearly delineate the mechanism in which DRP-104 may reduce T cell exhaustion or enhance T cell function. One hypothesis, referred to as “glutamine steal” phenomenon^86^, is that tumors with glutamine consumption can inhibit T cell function through depletion of free glutamine in the microenvironment. Potentially DRP-104 reverses this depletion by inhibiting glutamine consumption of *KEAP1* mutant tumors and thereby increasing extracellular glutamine availability for metabolically active effector T cells. While this hypothesis supports an indirect effect of DRP-104 on T cells, DRP-104 may also have a direct effect on T cells. Previous work has evaluated the effect of DON on T cells ^47^, but did not explore the effect of the prodrug DRP-104. It is also not clear to what extent CES1 is excreted and able to enzymatically activate DRP-104 in the extracellular space of the tumor microenvironment, where it can directly affect T cell function. Further work is needed to investigate the impact of DRP-104 on T cells in a reductionist manner, to specifically examine the impact on T cell activation, cytokine production, and exhaustion.

In summary, DRP-104 is a promising novel therapeutic agent that has high efficacy in *KEAP1* mutant lung tumors. Our work demonstrates that not only does DRP-104 target tumor intrinsic vulnerabilities via inhibition of nucleotide synthesis, but also enhances the function of anti-tumor T cells and can be combined with checkpoint blockade, the current standard of care in NSCLC. These findings provide a mechanistic rationale for the clinical trial (NCT04471415) using DRP-104 in combination with checkpoint blockade in LUAD patients specifically with loss-of-function *KEAP1* mutations or gain-of-function *NRF2* mutations.

## Methods

### Cell lines

KP and KPK cells utilized here were previously established^2^. *Stk11* knockout tumors were generated by transient transfection of PX458 (Addgene 48138) expressing a guide targeting *Lkb1*. Single GFP positive clones were selected and *Lkb1* loss was validated by western blot. *Nrf2* gain of function (Neh2 deletion), *Keap1* R470C mutant, *Keap1* WT KP cell lines were generated as previously described^2, 49^. Cells were cultured in DMEM with 10% FBS and gentamicin. *Keap1* R470C and *Keap1* WT KP cells were maintained in hygromycin selection (800 ug/mL).

### *In vitro* DRP-104 treatments and metabolic rescues

For cell viability assays cells were plated in a white, opaque 96-well plate with clear bottom at a density of 1000 cells/well in RPMI-1640 with 10% FBS. After attachment, DRP-104, CB-839, or DON were added at the indicated concentrations. After 5 days, cell viability was assessed by cell titer glo. For metabolic rescue experiments 2000 cells/well were plated in a 12 well plate in RPMI-1640 with 10% FBS. Cells were pre-treated with the indicated metabolites for 24 hours and then treated with DRP-104 for 5 days. Cells were stained with a 0.5% crystal violet solution in 20% methanol. Plates were then washed, dried, and crystal violet was eluted in 400mL of 10% acetic acid. Data is plotted as relative cell growth to vehicle treated control.

### Tumor mouse models

*In vivo* experiments using KP, KPK, KP *Nrf2* GoF, and *Keap1/Stk11* co-mutant KP cells were performed using nude (JAX strain# 002019), NOD SCID Gamma (NSG, JAX Strain #005557), of C57BL/6J (JAX strain #000664) mice. Cells (100,000 cells in 100 uL PBS) were injected subcutaneously into each flank of the mouse. Tumors were measured by calipers and volume was calculated based on 0.5 x length x width^2^. Once tumor volume was ∼ 100 mm^3^ treatment was initiated. For patient derived xenograft models (PDX), tumors were implanted in the flank of NSG mice as previously described^2^. To generate orthotopic lung tumors *Keap1* WT or *Keap1* R470C KP cells expressing luciferase GFP were injected intravenously (100,000 cells in 100 uL PBS) into female C57BL/6J (JAX strain #000664) mice and tumor burden was measured by bioluminescence (PerkinElmer IVIS Spectrum In Vivo Imaging System, D-luciferin PerkinElmer #122799). Data was analyzed using Living Image software.

### Treatments and T cell depletion

Treatment with DRP-104 (1 - 4 mg/kg) or vehicle control (10% tween 80, 10% ethanol in 0.9% saline) was administered subcutaneously five days on, two days off. For anti-PD1 therapy, mice were administered anti-PD1 (29.F1A12, BioXcell BE0273) or isotype control (rat IgG2a, BioXcell #BE0089) intraperitoneally 200 ug three times a week. For T cell depletion experiments either anti-CD8 (2.43, BioXcell BE0061) or isotype control (rat IgG2a, LTF-2, BioXcell BE0090) was administered 150 ug intraperitoneally twice a week once tumor burden was established by bioluminescence. One day after administration of antibody DRP-104 (3 mg/kg) was injected subcutaneously five days on, two days off.

### Metabolomics

For *in vitro* tracing, KPK tumor cells were plated at 100,000 cells per well in a 12 well plate in RPMI-1640 with 10% FBS. Cells were treated with DRP-104 (0.5 – 1.0 uM or DMSO) for 24 hours. Media was then replaced with fresh RPMI containing either 11 mM [U^13^C]-D-glucose or 2 mM [U^13^C]-L-glutamine and cultured for 1 hour. Cells were collected and prepared for LCMS as previously described^87^. For *in vivo* metabolomics, CTG743 PDX tumors were implanted into NSG mice as described above. After tumors were approximately 100 mm^3^ mice were treated with either vehicle control or DRP-104 (3 mg/kg) for five days. Mice were euthanized and then tumors were dissected. Tumor tissue was immediately flash frozen in liquid nitrogen. Approximately 5 mg of tissue was collected for analysis by LCMS. Tumor tissue was homogenized in metabolite extraction buffer (80% v/v ice cold methanol containing 1.4ug/mL norvaline) using a Precellys. After homogenization, tissue samples were prepared for LCMS following the same methods used for the *in vitro* tracing experiments described above. Fractional enrichment was calculated as the peak area of an individual isotopologue divided by the summed peak areas of all isotopologues for that metabolite.

### ExCITE seq

Mice were sedated with ketamine and xylazine and then were injected with 2 ug of APC anti-CD45 (2 ug per mouse diluted in 100 uL PBS, Biolegend 30-F11) retro-orbitally. After three minutes the chest of the mouse was opened. Lungs were removed and each lobe was separated and cleaned. The lung lobes were cut on a glass slide into small pieces followed and then digested (collagenase IV (Sigma Aldrich C5138), DNAse I (Sigma Aldrich, DN25) in RPMI with 10% FBS) for 35 minutes at 37 degrees. Digestion was stopped by addition of EDTA (1 mM). Digested cells were then filtered into a single cell suspension through a 100 micron filter. RBC lysis was performed. Cells were then washed and suspended in a staining buffer. Cells were then stained with live dead staining (Zombie UV fixable viability dye, Biolegend #423107) and PeCy7 anti-CD45 (see flow cytometry methods section for staining protocol).

Approximately 500,000 lung immune cells from each condition (2 mice per condition) were sorted as live^+^ IV-CD45 ^-^ CD45^+^ and 50,000 tumor cells were sorted as live^+^ CD45^-^ GFP^+^. Sorted samples were multiplexed using cell hashing antibodies (Biolegend) and stained with ADTs (see antibody table). 25,000 cells from each treatment condition were pooled and loaded into 10X Chromium. Gene expression together with Hashtag oligo (HTO) libraries were processed using Cell Ranger (v5.0.0) in multi-mode. Cell-containing droplets were selected using HTODemux function available in Seurat program. UMI count matrices from each modality were imported into the same Seurat object as separate assays. Viable cells were filtered based on having more than 200 genes detected and less than 10% of total UMIs stemming from mitochondrial transcripts. HTO counts were normalized using centered log ratio transformation before hashed samples were demultiplexing using the Seurat::HTODemux function. Protein counts were normalized using centered log ratio transformation. RNA counts were normalized using Seurat::SCTransform function with regressions of cell cycle score, ribosomal and mitochondrial percentages. Multimodal integration was performed using the weighted-nearest neighbor (WNN) method in Seurat. Briefly, a WNN network was constructed based on modality weights estimated for each cell using Seurat::FindMultiModalNeighbors function with top 40 and top 30 PCs from normalized RNA and protein counts, respectively. A shared nearest neighbor graph was then built based on the first 40 principal components (PCs) followed by identification of cell clusters using Leiden algoithm and Seurat::FindClusters function at multiple resolutions in order to identify potential rare cell types. Cell types were annotated based on canonical cell type markers and differential expressed genes of each cluster identified using Seurat::FindAllMarkers function with a logistic regression model. Clusters expressing markers of the same cell type were merged into a single cluster. Cell were then projected on to a uniform manifold^88^ using the top 40 PCs for visualization.

### Flow Cytometry

Mice were euthanized and lungs were digested into a single cell suspension as described above. Single cells were transferred to a 96 well round bottom plate and resuspended in FACS buffer (0.5% BSA, 0.1% sodium azide, and 1 mM EDTA). Live/dead staining was initially performed per protocol (Zombie UV fixable viability dye, Biolegend 423107). Cells were then blocked with Fc block (2.4G2, BioXcell) for ten minutes on ice. Antibody cocktail for surface staining was then added for 15 minutes on ice and then samples were washed with FACS buffer. Cells needing intracellular staining for FoxP3 were fixed and permeabilized using the FoxP3 Staining buffer kit (eBioscience 00552300). Intracellular Fc blocking was applied for 10 minutes on ice and then intracellularly stained with FoxP3 antibody for 1 hour on ice. Cells were then washed and resuspended in FACS buffer. For cytokine staining, single cell suspension cells were plated on a 96 well flat bottom plate. Cells were stimulated with PMA (0.1 ug/mL, Sigma P-8139), ionomycin (1ug/mL, Sigma I-0634), Golgi Plug (BD Biosciences 55029, 1:1000), and Golgi Stop (BD biosciences 555029, 1:1000) for 4.5 hours in RPMI with 10% FBS at 37°C. Next cells were washed, transferred to a 96 well round bottom plate and resuspended in FACS buffer. Surface staining was done as described above. Cells were then fixed with 2% PFA (diluted in FACS buffer) and then permeabilized by 0.5% saponin (diluted in FACS buffer). Cells were blocked intracellularly with Fc block and then stained for 1 hour with cytokine antibody cocktail. Next the cells were washed and resuspended in FACS buffer. The samples were filtered with a 100 micron filter and then run on BD LSRFortessa. Data was analyzed using FlowJo version 10. Antibodies used are listed in Supplementary Table 1.

### Statistics

Statistical analysis was performed using GraphPad Prism v9. All data is expressed as mean plus standard error of the mean. Data was analyzed by statistical test indicated in figure legends. All tests were two tailed.

### Data Availability

Data is available from the corresponding author upon request.

---

**Extended Data Figure 1:**
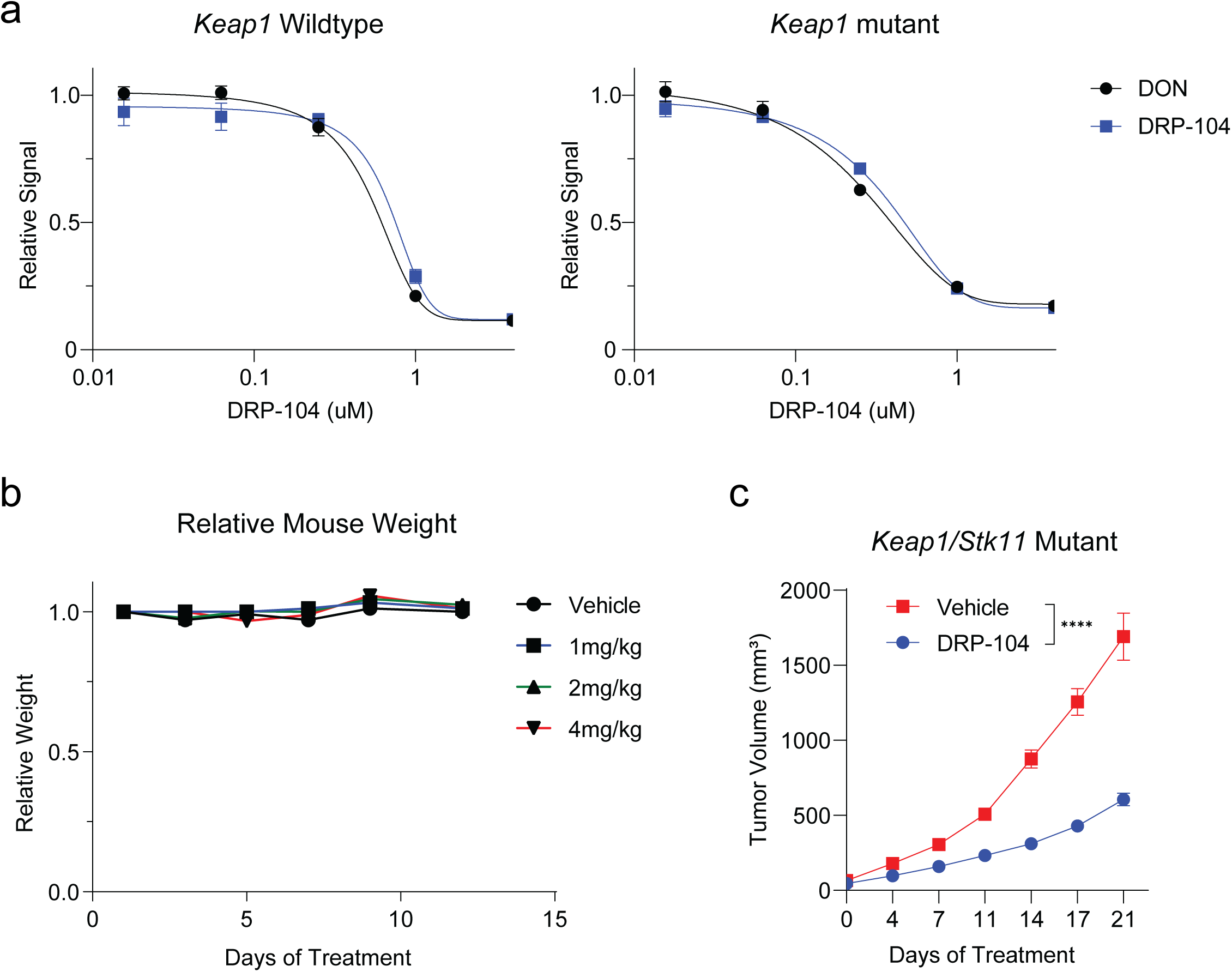
DRP-104 is effective *in vitro* and *in vivo* against *Keap1* mutant tumors. **a:** *In vitro* dose response to DRP-104 and DON in *Keap1* wildtype and mutant cells. Cells were plated and treated with drug for five days and viability was measured by cell titer glo. **b:** C57BL/6 mice were treated with DRP-104 1, 2, or 4 mg/kg and relative weight to baseline is plotted. **c:** *Keap1/Stk11* co-mutant *Kras*^G12D/+^ *p53*^-/-^ cells were transplanted subcutaneously into C57BL/6 mice and response to DRP-104 (3 mg/kg) was measured (n=10 per group). Tumor growth was analyzed by two-way ANOVA. **** p < 0.0001.

**Extended Data Figure 2:**
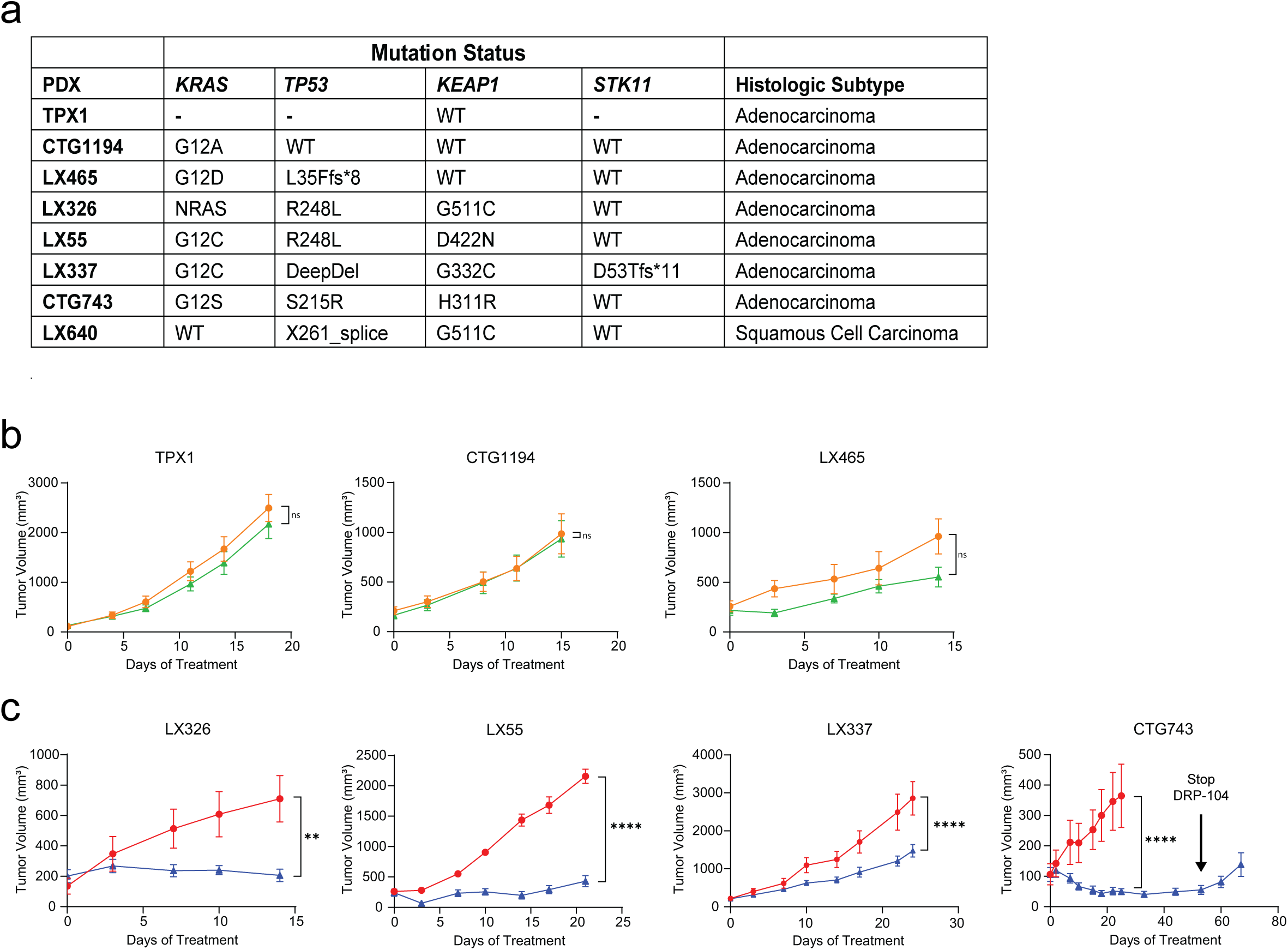
*KEAP1* mutant patient derived xenografts respond to DRP-104. **a:** Table with patient derived xenograft (PDX) lines with mutation status and histologic subtype listed. **b, c:** Treatment responses as measured by PDX tumor volume to DRP-104 (3 mg/kg) or vehicle control in **b**) *KEAP1* wildtype PDX (TPX1: n=8-10 per group, CTG1194: n=14-16 per group, LX465: n=8-9 per group) and **c**) *KEAP1* mutant PDX (LX326: n=6 per group, LX55: n=8-9 per group, LX337: n=8 per group, CTG743: n=10 per group). Timepoint of withdrawal of DRP-104 is indicated for CTG743 PDX experiment. Tumor growth was analyzed by two-way ANOVA. * p < 0.05, ** p<0.01, **** p < 0.0001.

**Extended Data Figure 3:**
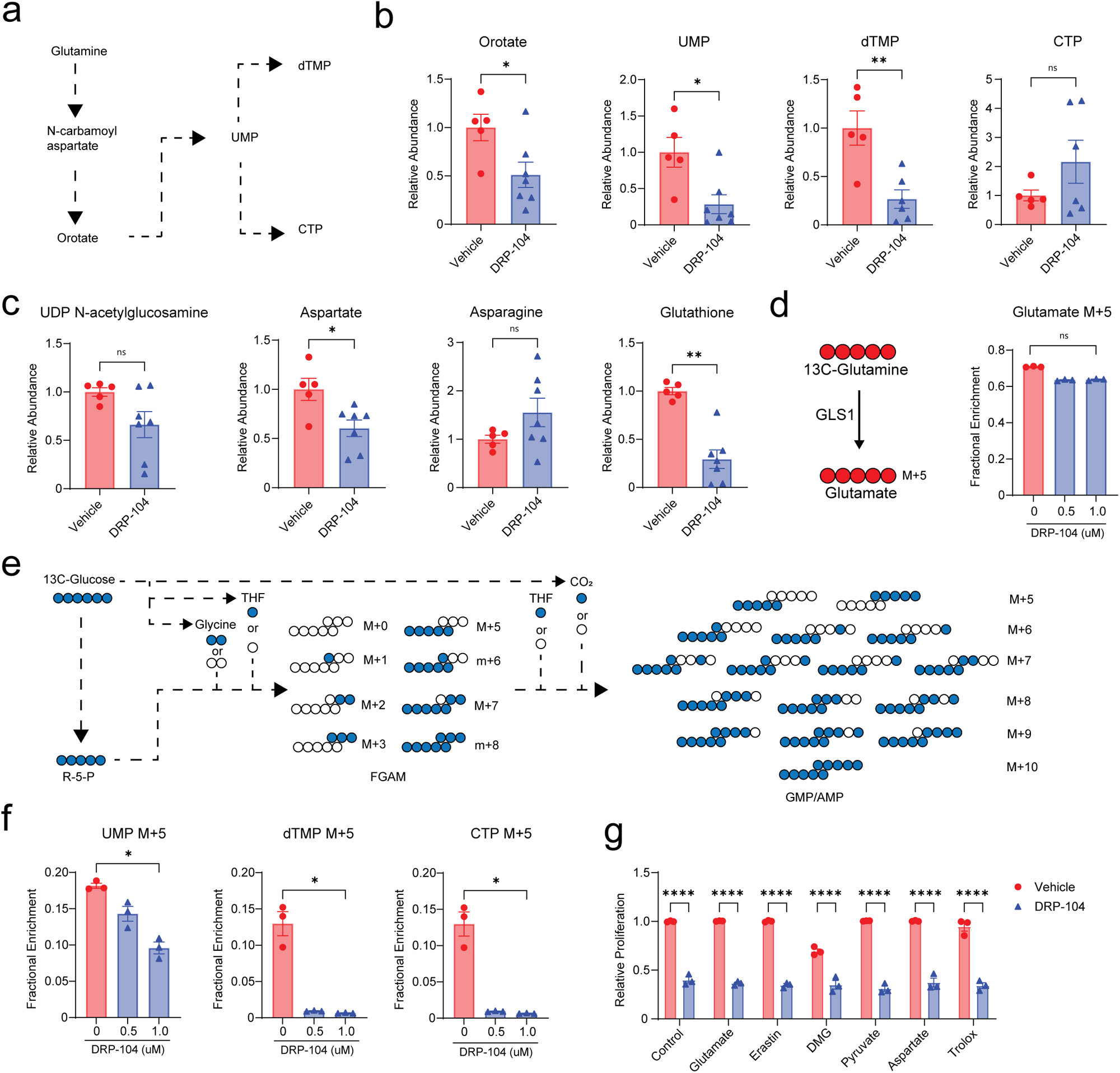
*In vivo* and *in vitro* metabolomics of DRP-104 treated *KEAP1* mutant tumors. **a:** Schematic of pyrimidine synthesis pathway. **b, c:** Levels of selected metabolites by liquid chromatography mass spectrometry in CTG743 (*Keap1* mutant) patient derived xenograft tumors after 5 days of DRP-104 treatment (n=5-7 per group). **d:** *In vitro* tracing of *Keap1* mutant tumor cells with DRP-104 (n=3 per group) after 1 hour of labeling with 13C-glutamine (left) and quantification of fractional enrichment of 13C-glutamine labeled glutamate for each concentration of DRP-104 (right). **e:** Schematic of 13C-Glucose tracing for purine synthesis. **f:** Fractional enrichment of pyrimidines after 13C-Glucose labelling with DRP-104 treatment *in vitro* (n=3 per condition). **g:** *Keap1* mutant tumor cells were pretreated with specific metabolites 24 hours prior to DRP-104 treatment (2 uM). After five days of drug or vehicle treatment, proliferation was measured by crystal violet and plotted relative to controls (n=3 per condition). Statistical analysis was done by either Mann Whitney test, Kruskal-Wallis test with Dunn’s multiple comparisons test, or 2-way ANOVA. ns: not significant, * p < 0.05, ** p<0.01, **** p < 0.0001.

**Extended Data Figure 4:**
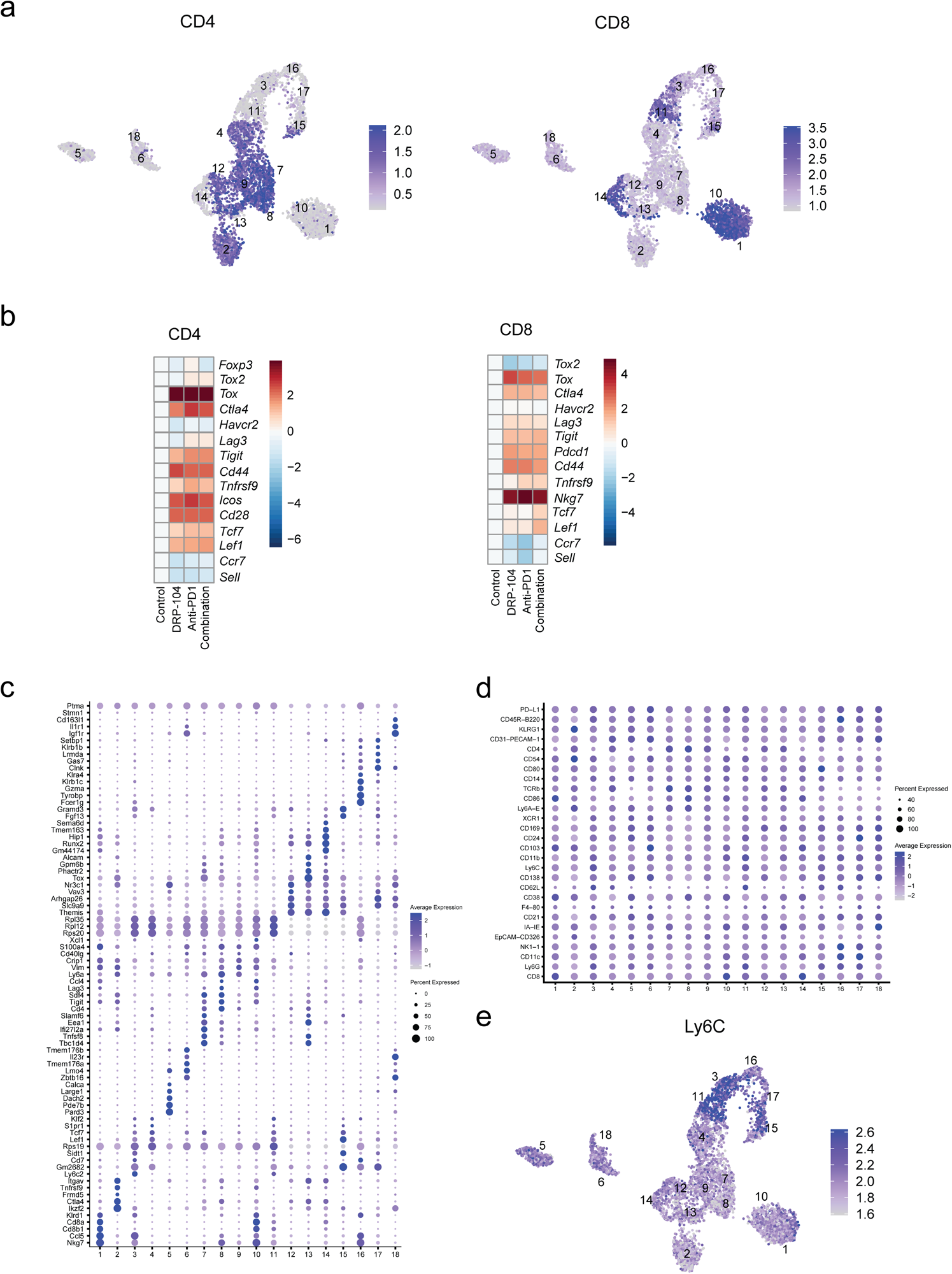
ExCITE-seq Analysis of T/NK/NKT/ILC Cells. **a:** Antibody derived tag (ADT) expression for CD4 and CD8 on T cell/NK cell/NKT cell/ILC subclusters. **b:** Heatmap for expression of selected genes in CD4 and CD8 T cells. **c:** Differentially expressed genes for T/NK/NKT/ILC subclusters. **d:** ADT expression by T cell/NK cell/NKT cell/ILC subcluster. **e:** Protein expression of Ly6C by ADT.

**Extended Data Figure 5:**
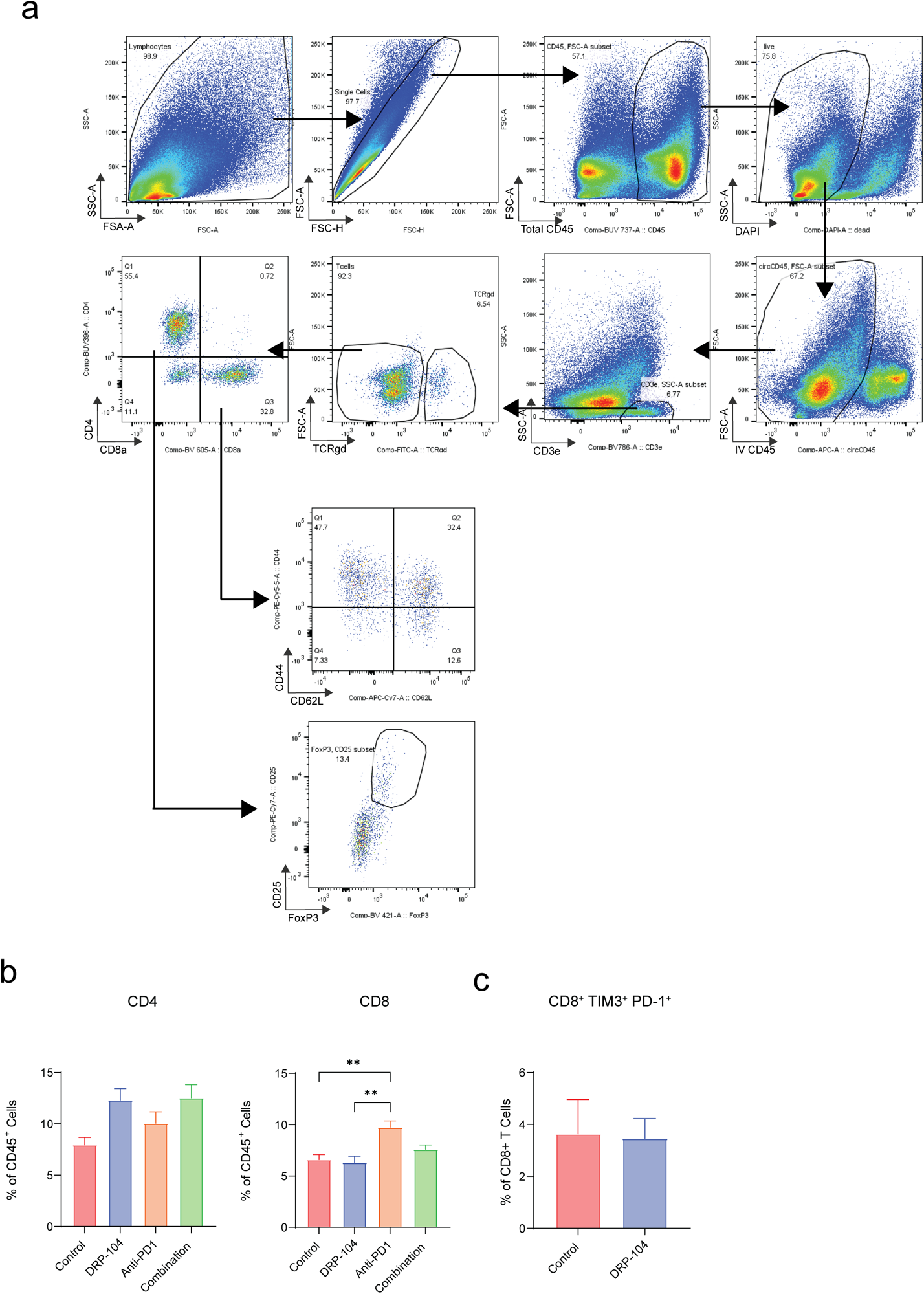
Flow Cytometry Analysis of T cells from orthotopic lung cancer model. **a:** Gating strategy for T cell populations. **b, c:** Flow cytometry quantification of (**b**) CD4, CD8, and (**c**) CD8^+^ TIM3^+^ PD-1^+^ T cell populations of *Keap1* R470C mutant tumor bearing lungs after five days of DRP-104 (3 mg/kg daily) or vehicle and anti-PD1 (200 ug three times a week) or isotype control treatment (n=5 per group). Statistical analysis was done by one-way ANOVA with Tukey’s multiple comparisons test or Mann Whitney test. ** p<0.01

Supplementary Table 1.

**Supplementary Table 1**
| Antigen | Fluorophore | Clone | Manufacturer | Catalogue # |
| --- | --- | --- | --- | --- |
| CD45 | APC | 30-F11 | Biolegend | 103112 |
| CD45 | BUV737 | 30-F11 | BD Biosciences | 748371 |
| CD3e | BV785 | 17A2 | Biolegend | 100232 |
| CD4 | BV395 | GL15 | BD Biosciences | 565974 |
| CD4 | Pacific blue | RM4-5 | Invitrogen | MCD0428 |
| CD8a | BV605 | 53-6.7 | Biolegend | 563152 |
| CD8a | PerCPCy5.5 | 53-6.7 | BD Biosciences | 45-0081-82 |
| TCRgd | FITC | UC7-13D5 | Biolegend | 107512 |
| CD62L | biotin | MEL-14 | eBioscience | 13-0621-85 |
| CD44 | PerCPCy5.5 | IM7 | eBioscience | 45-0441-80 |
| FoxP3 | e450 | FJK-16S | eBioscience | 48-5773-82 |
| LAG3 | PeCy7 | C9B7W | eBioscience | 25-2231-80 |
| TIM3 | PE | RMT3-23 | Biolegend | 119703 |
| TCF7 | AF488 | C63D9 | Cell Signaling Technologies | 6444S |
| IFNγ | FITC | XMG1.2 | BD Biosciences | 557724 |
| TNFα | PeCy7 | MP6-XT22 | eBioscience | 25-7321-82 |
| PD-1 | BV421 | RMP1-30 | eBioscience | 109121 |
| CD25 | PeCy7 | PC61.5 | eBioscience | 25-0251-82 |

## Acknowledgements

R. P. was supported by the William Rom fellowship, the Stony Wold-Herbert Fund, and NIH training grant T32 CA009161 and T32 AI100853. T. P. is supported by NIH grants (R37CA222504 and R01CA227649) and an American Cancer Society Research Scholar Grant (RSG-17-200-01–TBE). Work in S.B.K. laboratory was supported by NIH (R01HL-125816, R01CA271245), LEO Foundation Grant (LF-OC-20-000351), NYU Cancer Center Pilot grant (P30CA016087). PDX generation supported by NIH P30 CA008748 and the Druckenmiller Center for Lung Cancer Research. The study was partly supported by Dracen Pharmaceuticals Inc. We thank members of the Experimental Pathology Research Laboratory, which is partially supported by the Cancer Center Support Grant P30CA016087 at NYU Langone’s Laura and Isaac Perlmutter Cancer Center. The Akoya Vectra Polaris multispectral scanning system was awarded through the shared instrument grant S10 OD021747.

## Authors Contributions

R.P., S.E.L., S.B.K., and T.P. conceived the project, designed the experiments, and wrote the manuscript. R.P., S.E.L., A.R., S.H.M., C.B., W.L.W, B.K., A.H., E.I., M.C., J.B., M.H., S.R., V.I.S., and T.K. performed experiments. R.P., S.E.L., analyzed *in vitro* and *in vivo* data. C.N. and J.B. analyzed the metabolomics data. Y.H. performed the single cell analysis. V.I.S., K.M.K., and K.K.W. provided conceptual advice. R.W. provided conceptual advice and edited the manuscript. A.T. supervised the single cell analysis. J.T.P. and C.M.R. provided PDX models. S.M.D. supervised the metabolomics experiments/analysis. S.B.K. supervised the immune analysis and single cell analysis. T.P. supervised the study. All authors reviewed the manuscript.

